# FrenchFISH: Poisson models for quantifying DNA copy-number from fluorescence *in situ* hybridisation of tissue sections

**DOI:** 10.1101/487926

**Authors:** Geoff Macintyre, Anna M. Piskorz, Edith Ross, David B. Morse, Ke Yuan, Darren Ennis, Jeremy A. Pike, Teodora Goranova, Iain A. McNeish, James D. Brenton, Florian Markowetz

## Abstract

Chromosomal aberration and DNA copy number change are robust hallmarks of cancer. Imaging of spots generated using fluorescence *in situ* hybridisation (FISH) of locus specific probes is routinely used to detect copy number changes in tumour nuclei. However, it often does not perform well on solid tumour tissue sections, where partially represented or overlapping nuclei are common. To overcome these challenges, we have developed a computational approach called FrenchFISH, which comprises a nuclear volume correction method coupled with two types of Poisson models: either a Poisson model for improved manual spot counting without the need for control probes; or a homogenous Poisson Point Process model for automated spot counting. We benchmarked the performance of FrenchFISH against previous approaches in a controlled simulation scenario and exemplify its use in 12 ovarian cancer FFPE-tissue sections, for which we assess copy number alterations in three loci (c-Myc, hTERC and SE7). We show that FrenchFISH outperforms standard spot counting approaches and that the automated spot counting is significantly faster than manual without loss of performance. FrenchFISH is a general approach that can be used to enhance clinical diagnosis on sections of any tissue.

**Author summary:** Cancer genomes can look very chaotic, because cancer cells are unable to fully repair errors in DNA replication during cell division. While a healthy genome has two copies of every chromosome, in a cancer genome some pieces can be lost completely and others can appear in 50 copies. To diagnose cancers and to decide on the right therapeutic strategy for a patient, it can be very important to know how many copies of a particular piece of DNA exist in a cell. The standard technique used in the clinic to assess DNA copy number is called FISH, short for fluorescence *in situ* hybridisation. This technique uses fluorescent probes that bind to a DNA piece of interest and show up as glowing spots in a microscopic image. Counting the spots in an image is a labour-and time-intensive process that is generally done by well-trained experts. Here we present a statistical approach to automatically count FISH spots, which outperforms previously proposed methods, and has the potential to substantially speed up clinical diagnostics.

## 1 Introduction

Chromosomal instability coupled with defective DNA repair can cause loss or duplication of DNA, a characteristic attribute of cancer cells [1]. Interrogation of DNA copy number aberrations is critical for diagnosis [2] and understanding tumour etiology [1]. Technologies for measuring DNA copy-number have evolved from optical profiling of single loci [3] through to sequencing of the entire tumour genome [4]. However, determining the absolute number of copies from bulk sequencing data remains difficult because of normal cell contamination and intra-tumour heterogeneity [5], and results are generally reported in terms of loss or gain of DNA relative to an assumed diploid or median background. Information from single locus methods is therefore often required to validate estimates of absolute copy-number [6, 7].

Fluorescence *in situ* hybridisation (FISH) of interphase nuclei is the most widely established technique for interrogating single locus copy number. Fluorescent probes are hybridised to a specific genomic region of interest and appear as discrete foci when visualised with fluorescent microscopy [8]. Standard analysis of FISH data relies on time-consuming manual counting of spots in these images [9]. Automated systems to quantify foci using nuclei recognition and spot counting algorithms (reviewed in [10]) aim to make the analysis of FISH data less labour-intensive, faster, and more objective. However, the accuracy of most systems is limited to identification of spots in intact and separated nuclei [11]. Thus, these systems can be very successful in haematological malignancies; however, diagnostic sections of solid tumour tissue pose a significant challenge for both automated and manual analysis. Accurate identification of single nuclei either by eye or by automatic image segmentation can be hard if nuclei cluster closely and overlap (see Fig 1). Arbitrary cut points between grouped nuclei are typically used to separate these clusters, which can lead to noisy spot count estimates. Additionally, tissue sections are typically 3 μm to 5 μm, which is smaller than the diameter of most tumour nuclei, and thus the majority of nuclei are not captured completely in the volume of the section [12, 13]. To address these challenges, both manual and automated analysis have been improved by using control probes that bind to a specific locus with known copy-number state *n*_ctrl_ [10]. Two commonly used approaches are:

1. Only nuclei containing the expected number of control probes (usually *n*_ctrl_ = 2) are used to estimate the copy-number of other loci. The underlying assumption is that if a nucleus contains the expected number of control probes then it is likely that the majority of the nucleus is captured by the section and hence other spots will be well represented.
2. The spot count for the locus of interest is scaled by the ratio of expected over observed control probe copy-number:

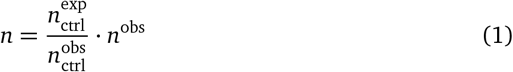

**Fig 1.**
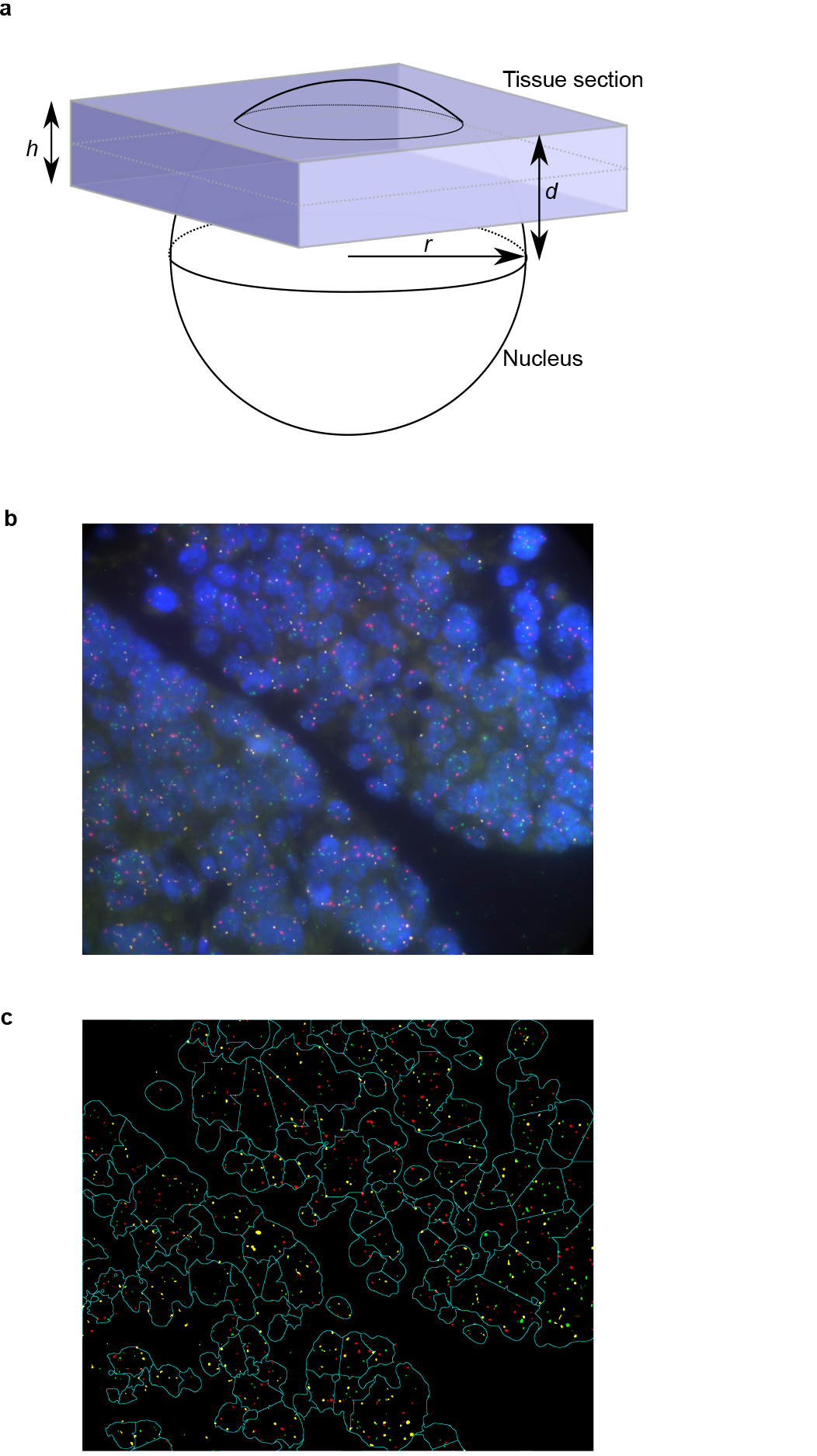
Tissue section fish. a: A schematic of a tissue section cutting through a cell nucleus. The highlighted quantities are used for calculating the volume of nucleus appearing in the tissue section. b: Three probe fish applied to high-grade serous ovarian cancer. c: Automatic image segmentation and spot recognition applied to the image in b. Notice the difficulty in accurately separating overlapping nuclei.

In this case, the underlying assumption is that the number of observed control spots is linearly correlated with the number of spots observed for the locus of interest.

However, there are significant limitations associated with both of these methods. For example, a tissue section 3 μm thick containing cells with a nuclear diameter of 9 μm will, on average, have only 41% of each nucleus represented in the section (see Fig 1a). Therefore, for method 1, it is unlikely that the section will contain many nuclei with a complete control probe count and the locus of interest is likely to be undersampled. Using thicker tissue sections can overcome this limitation. However, as section thickness increases, the quality of imaging decreases and many more overlapping nuclei are captured, which complicates identification of single nuclei. Method 2 performs well when the control probe is at the expected copy-number. However, in tumours with significant aneuploidy, it is difficult to identify a control probe with constant copy-number, even when using centromeric probes.

Thus, new automated approaches are required that generate robust and reproducible results from fixed tumour sections. Ideally, new methods should account for the three major challenges in FISH analysis of tissue sections: (1) nucleus subsampling, (2) control probe aneuploidy and (3) overlapping nuclei. We have addressed these challenges by developing FrenchFISH, a computational package that comprises three major computational innovations for improved spot counting: volume adjusted spot counting, which accounts for partial nucleus representation without the need for control probes; Poisson estimated spot counts from manually counted nuclei, which account for uncertainty in spot counts; and a homogeneous Poisson point process model, which facilitates automated spot counting and circumvents the need for single nucleus image segmentation. In the following, we present the details of the FrenchFISH model, show that it outperforms standard spot counting approaches and is significantly faster than manual spot counting.

## 2 Results

FrenchFISH is implemented in R [14] utilising elements of the FishalyzeR [15] package on Bioconductor [16]. Scripts to reproduce all results are available as part of the supplementary information and can be found in the following repository: https://bitbucket.org/britroc/frenchfish

### 2.1 The FrenchFISH model

The goal of the analysis is to estimate the copy-number of a locus denoted by *n*, which we will achieve by volume-adjusting observed spot counts and using Poisson models.

#### 2.1.1 Observed spot counts

FISH of a probe specific to the locus allows us to observe copy-number in terms of spot counts inside the nucleus. Here, we assume a FISH image has 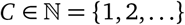 cell nuclei and the number of observed spots in cell 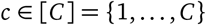 is 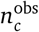. The average number of observed copies of the locus in the tissue section is

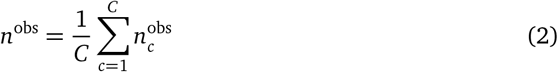

#### 2.1.2 Volume adjusted spot counting

Figure 1a displays a schematic of a nucleus subsampled due to tissue section cutting. For simplicity, we assume that all nuclei are spherical with radius *r* (*r* is typically estimated from image). Their volume then is given by

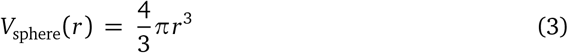

For a specified section thickness *h*, we can express the volume of the nucleus sampled by a section in terms of *d*, the distance of the section edge from the nucleus midline:

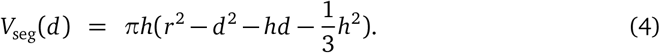

By integrating over *d* and dividing by *h*, we can compute the average volume sampled:

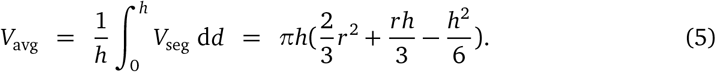

This quantity can be used to scale the observed number of spots to get an estimate of the true number of spots:

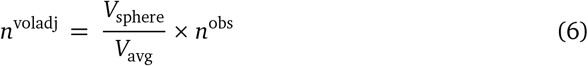

#### 2.1.3 Modelling uncertainty in manual spot counts

As the observed spot counts are subject to both hybridisation and image signal processing noise, we use a probabilistic model that accounts for this uncertainty. We model the counts as coming from a Poisson distribution with rate λ. Given this, the likelihood of our data can be expressed as

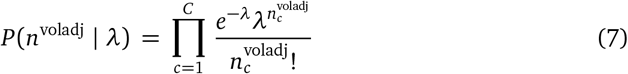

To compute the posterior of λ given the data, we use Bayes’ rule to transform the likelihood into

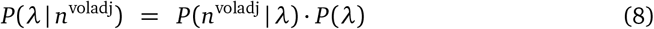

Using the conjugate Gamma prior as *P*(λ) and the likelihood of Eq. 7, we sample from the posterior with Markov Chain Monte Carlo (MCMC) to generate λ_*t*_ ∈ [*T*] values fit to the data after a burn-in of 1000 iterations. We use the `MCpoissongamma` function from the MCMCpack package [17] in R to achieve this. From this sampling chain we then compute the expected rate which is equal to the expected spot count:

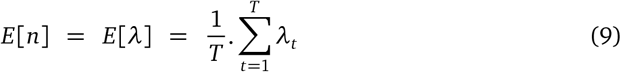

#### 2.1.4 Modelling uncertainty in automatic nuclear segmentation

While segmentation of single nuclei in tumour sections is difficult, separating nuclear staining from background and accurately defining spots remains relatively easy. Our approach exploits this fact in the framework of a Homogeneous Poisson Point Process. A Poisson Process models a continuous series of events across space or time. In our setting, we consider spots as events and nuclear area *a* measured in *μm*^2^ as space. The number of spots in an area *a* is denoted by *N*(*a*) and modelled by a Poisson process with intensity λ^PP^:

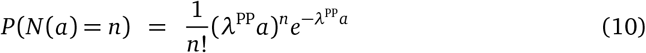

and using the `fitPP.fun` from the NHPoisson package [18] in R, we obtain a maximum likelihood estimate for λ^PP^.

As λ^PP^ is a spot count estimate per *μm*^2^ of observed nuclea area, to get the estimated number of spots per nucleus, we first multiply by the average area of a nucleus, *πr*^2^, and then scale by the average nuclear volume represented in the tissue section, to get an estimate of the number of copies *n*:

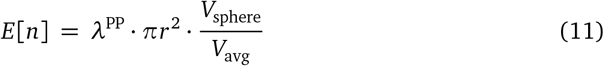

### 2.2 Validation and benchmarking of FrenchFISH

To validate and benchmark FrenchFISH, we used the controlled scenario of a simulation study as well as a real-world case study in ovarian cancer.

#### 2.2.1 Benchmarking in simulation study

We simulated a total of 11,200 tissue sections to benchmark our approach. For each condition, we simulated 10 replicate sections with 50 nuclei. All nuclei had their midpoint location randomly positioned within the tissue section. Test conditions were selected from all possible combinations of the following:

- control probe copy-number *n*_*control*_ ∈ {1, 2, 3, 4},
- probe of interest copy-number *n* ∈ {1, 2, 3, 4, 5, 6, 7, 8, 9, 10},
- percentage of nuclei with a probe sampling error of minus/plus one count *e* ∈ {−20, −10, −5, 0, 5, 10, 20},
- probability of nucleus overlapping with another nucleus in the section *p* ∈ {0, 10, 30, 50, 80}.

Using these data we tested FrenchFISH’s performance against the standard approach outlined in equation 1 where a control probe (assumed to be diploid) is used to scale the observed spot counts.

##### Benchmarking against noisy spot counts

We first measured performance using simulated tissue sections with non-overlapping nuclei and varying levels of noise. Noise was introduced by either under counting or over counting by one spot, in 5,10 or 20% of the cells in each tissue section.

Naive spot counting without correction showed a severe underestimate of the true number of spots (Figure 2a). The standard correction approach improved spot count estimates when the control probe was diploid (Figure 2b). However, estimates showed high variability as the true number of spots increased. In contrast, FrenchFISH showed consistent performance across all true copy number states. Performance remained adequate for noise levels up to 10%. The standard approach showed worse performance than FrenchFISH at 20% noise, however, errors were less pronounced for under counting noise compared to over counting noise (Figure 2b). The standard approach largely failed to provide correct copy number estimates when the control probe copy number was other than diploid, especially for higher noise levels (Figure 2c). FrenchFISH did not show a deterioration in performance as it did not rely on a control probe.

**Fig 2.**
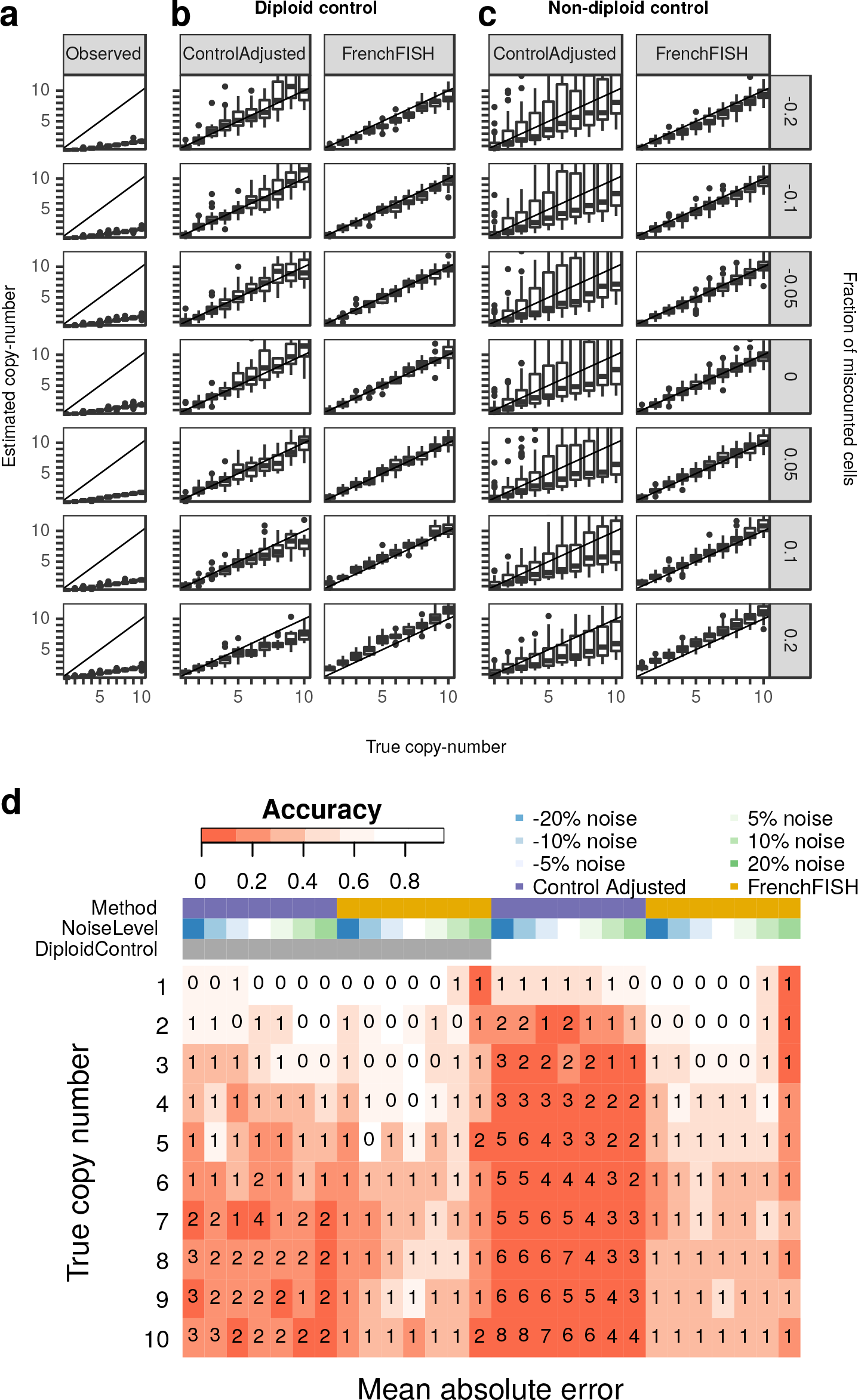
Performance assessment on simulated spot counts. a: Box plots showing the distribution of unadjusted observed spot counts (y-axis) compared to the true spot counts (x-axis), for varying noise levels (y facets). b: Spot count estimates for the standard control probe adjusted method (Control Adjusted) and FrenchFISH. All simulated tissue sections in these plots had an accompanying diploid control probe count. c: Spot count estimates for tissue sections with non-diploid controls. d: A heatmap showing the accuracy of spot counting (shading) for each method, noise level, and control probe count. The integers inside the heatmap boxes show the mean absolute error.

To gain further insight, we observed accuracy and mean absolute error for both approaches under the same varying noise conditions (Figure 2d). Overall accuracy was poor for the standard approach except when the control probe was diploid and true copy number was 1. High accuracy was observed for FrenchFISH up to a true spot count of 4 and noise levels of 10%. Accuracy was poor in cases where over counting noise was 20%. Despite a deterioration in accuracy beyond true copy number counts of 4, FrechFISH’s mean absolute error never exceeded 1, thus FrenchFISH’s estimates were only ever wrong by one copy. In contrast, the standard approach had a mean absolute error of up to 7 under some conditions.

##### Benchmarking against overlapping nuclei

Here we assessed the performance of both methods across simulated tissue sections with varying degrees of nuclear overlap (Figure 3). Both methods were robust to nuclear overlap in the diploid control probe setting, including at 80% probability of overlap. However, the standard approach again showed more variable results as the true copy number increased (Figure 3b). The standard approach consistently failed to estimate the correct copy number when the control probe was not diploid, however, this error did not vary with degree of overlap (Figure 3c). FrenchFISH showed a mean absolute error no greater than one, whereas the standard approach showed up to 2 copies in the diploid control probe setting and up to 7 copies in the non-diploid setting (Figure 3d).

**Fig 3.**
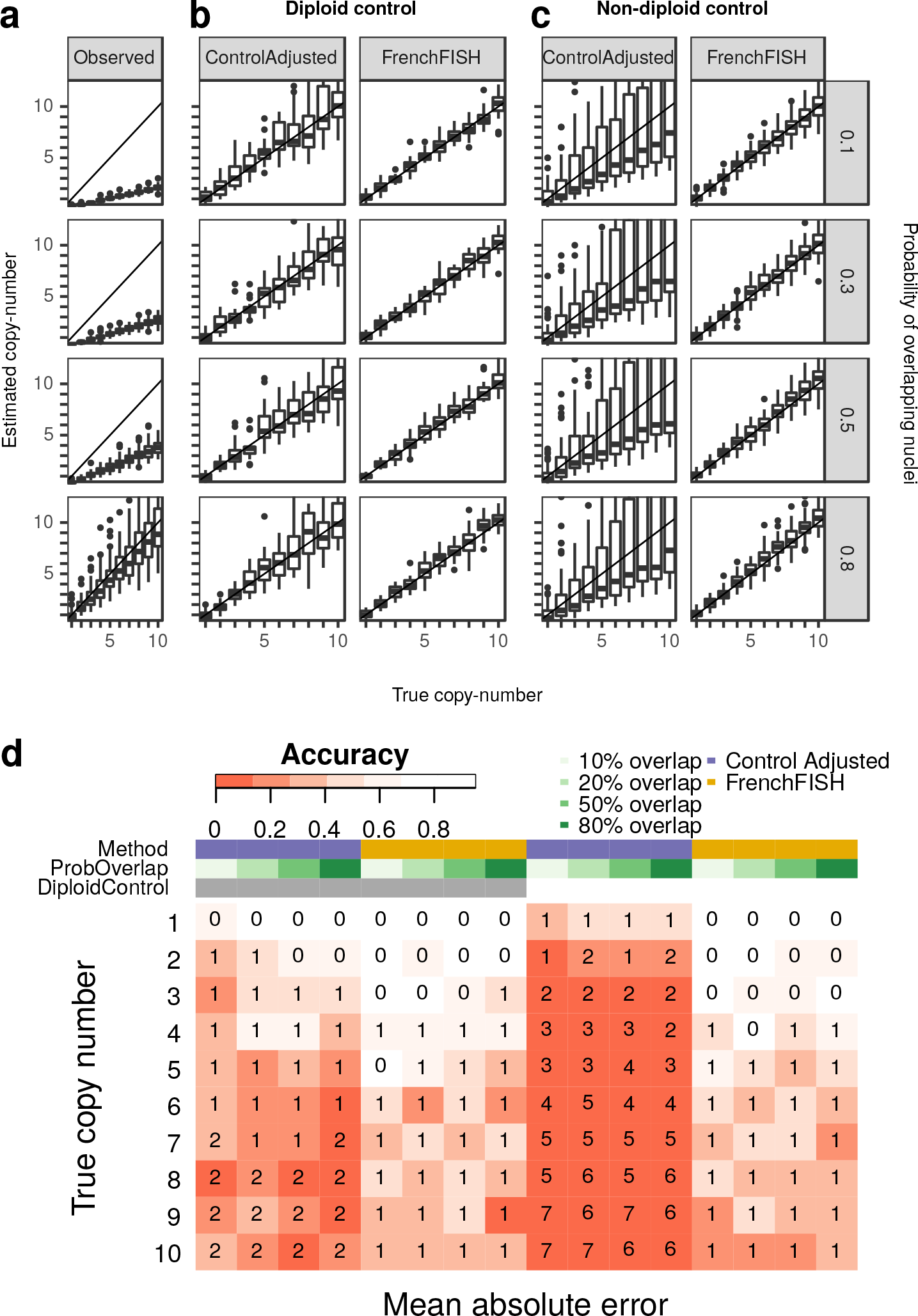
Performance assessment on simulated spot counts. a: Box plots showing the distribution of unadjusted observed spot counts (y-axis) compared to the true spot counts (x-axis), for varying levels of probability of overlapping nuclei (y facets). b: Spot count estimates for the standard control probe adjusted method (ControlAdjusted) and FrenchFISH. All simulated tissue sections in these plots had an accompanying diploid control probe count. c: Spot count estimates for tissue sections with non-diploid controls. d: A heatmap showing the accuracy of spot counting (shading) for each method, overlap probability, and control probe count. The integers inside the heatmap boxes show the mean absolute error.

#### 2.2.2 Case study on ovarian cancer tissue sections

We performed both manual and automatic spot counting on multichannel FISH of tissue sections from 12 ovarian cancer cases. Manual spot counts were corrected using FrenchFISH’s volume adjustment method and automatic counting was performed using the Poisson point process model.

##### Manual versus automatic counting

We observed the degree of agreement between manual and automatic spot counting to assess whether the automatic method resulted in any loss of performance compared to manual counting. 74% (26 of 35) of the estimated copy number counts were less than one copy number of each other with a further 17% (6 of 35) having estimates less than two copies different (Figure 5).

##### Timing analysis

We measured the time it took to perform both manual and automatic spot counting. Figure 4 provides a breakdown of the two approaches and the timings associated with each step. Using up to 5 fields of view per sample we were able to obtain roughly 100 manually curated nuclei per sample. The total average processing time for the automatic FrenchFISH approach was 36 minutes, 30 minutes for manual estimation of the nuclear diameter then 6 minutes software processing. The total average processing time for the manual approach was 119 minutes, with the majority of processing performed by a human.

**Fig 4.**
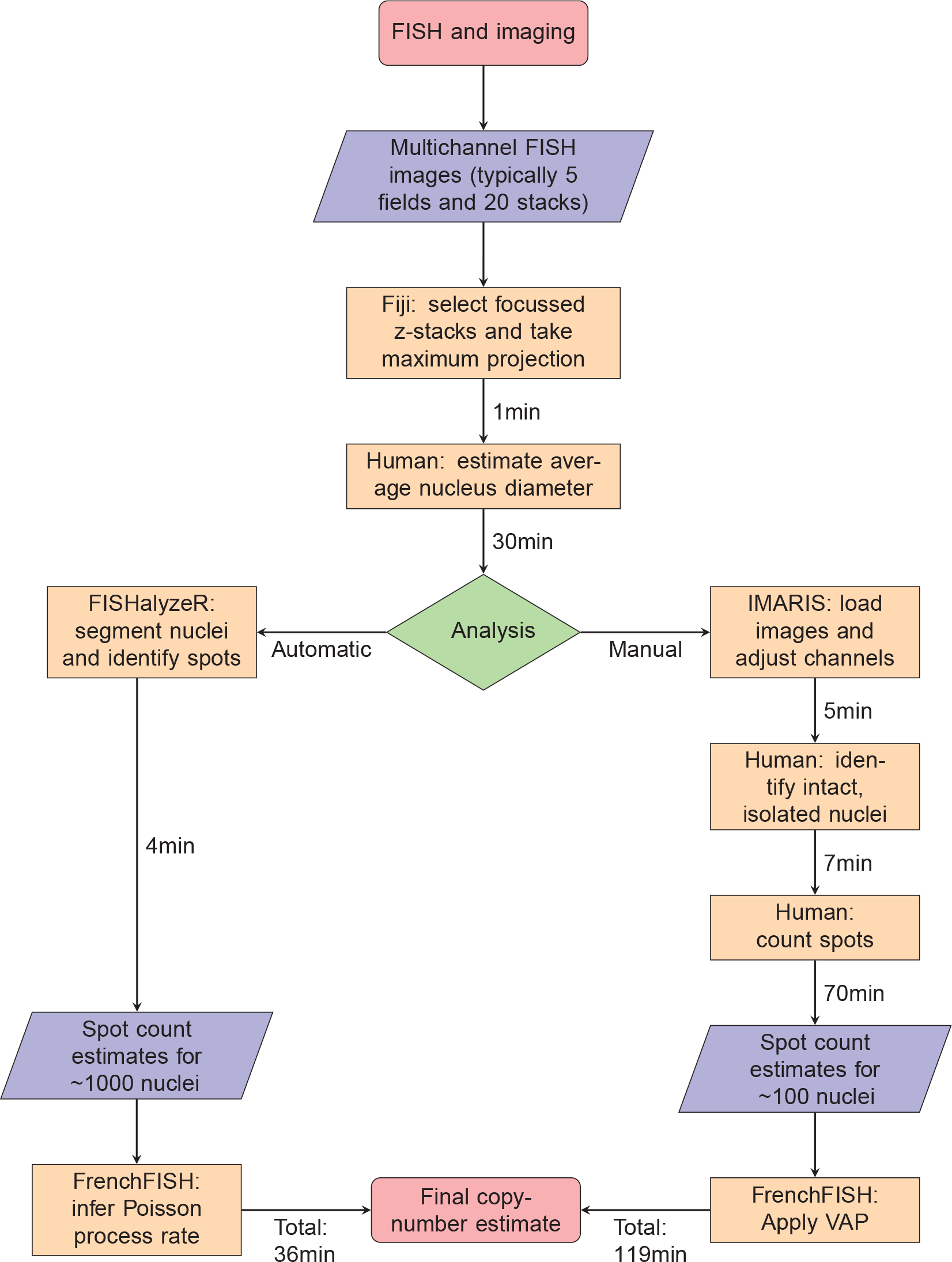
Comparison of manual spot counting and FrenchFISH’s automated spot counting, across 3 probes, using FISH of 12 ovarian cancer cases. This flow chart outlines the tasks required to carry out automatic or manual spot counting for a single sample. The minutes associated with each process are an average across 8 cases for up to 5 fields of view. Squares represent processes, the diamond represents a decision point and the trapezoids represent input/output. For each process it is listed whether it is carried out by software, or by human.

**Fig 5.**
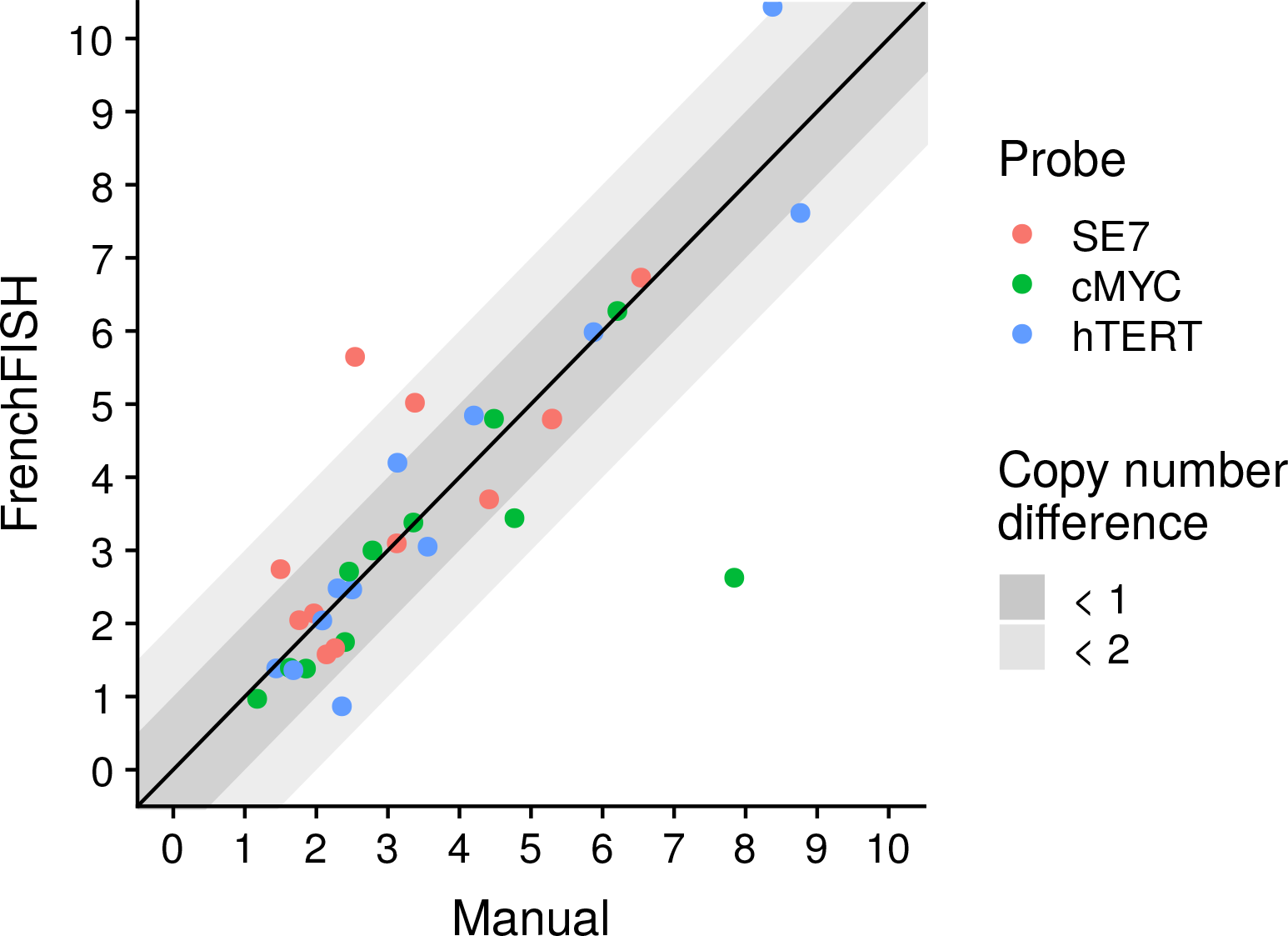
Spot count estimates for 3 probes across 12 ovarian cancer cases. This scatter plot shows spot count estimates from manual counting versus FrenchFISH’s automated spot counting. Points falling within the dark shaded area have estimates within 1 copy of each other across the methods. Those falling within the light grey area are within 2 copies. Those falling outside this area are greater than two copies different.

## 3 Discussion

Here we present FrenchFISH, a software tool for quantitative copy number estimation from FISH of tissue sections. We demonstrated the robust and superior performance of FrenchFISH using simulated tissue sections and FISH of ovarian cancer tissue sections. We explored the limitations of FrenchFISH using simulations of tissue sections with spot counting noise and overlapping nuclei. FrenchFISH was robust to overlapping nuclei noise and performed well in cases with up to 10% spot counting noise. Interestingly, over counting noise resulted in worse performance than under counting, suggesting that a conservative spot counting strategy could improve copy number estimates. Our controlled simulated setting also highlighted the difficulty in estimating high copy number states, with accuracy rapidly decreasing with copy numbers greater than 4 copies. However, in all cases tested, FrenchFISH’s estimates were not more than 1 copy different from the underlying truth.

On ovarian cancer tissue section, 74% of FrenchFISH’s automated spot count estimates were within 1 copy of manual counted estimates. This demonstrates that FrenchFISH is a viable alternative to manual counting, which would decrease analysis time fourfold with significantly less human intervention.

FrenchFISH is the first method specifically designed to provide quantitative copy number estimates from tissue section FISH without the need for a matched control probe.

## 4 Materials and methods

### 4.1 Simulation

Simulated tissue sections were generated using the following procedure:

1. Fix tissue section height *h* at 3μm and the nuclei radius *r* to 9μm
2. For *C* = 50 cells per simulated tissue section, estimate *d* (the distance from the midline of the nucleus to the top of the tissue section):

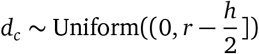
3. For each *d*_*c*_, calculate the fraction of the nucleus contained in the section,

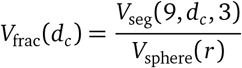
4. Using *V*_frac_ as the prior probability for seeing a spot sampled from a Poisson distribution, generate observed spot counts

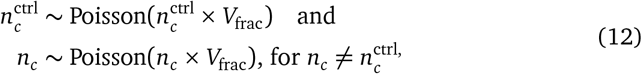
5. If the probability of overlap *p* is > 0, merge with neighbouring nucleus *c* + 1, recalculating the overlapped nuclei area *a*:
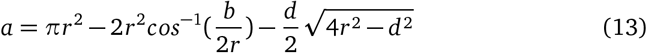

where *b* is the distance between nuclei centrepoints sampled from *d* ~ Uniform((0, 0.3]), and updated spot count:

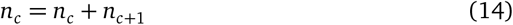
6. If error *e* is ≠ 0 then update spot count:

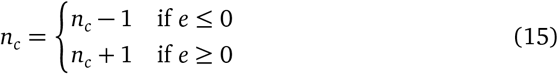
7. Repeat above steps 10 times for all possible combinations of *e* ∈ {−0.2, −0.1 −0.05, 0, 0.05, 1, 0.2}, *p* ∈ {0, 0.1, 0.3, 0.5, 0.8},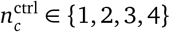 and *n*_*c*_ ∈ {1, 2, 3, 4, 5, 6, 7, 8, 9, 10}.

### 4.2 FISH on ovarian cancer tissue sections

#### 4.2.1 Patient sample selection

Eight high-grade serous ovarian cancer samples were selected and reviewed by a pathologist who marked the area of each tumour on the H&E sections. In addition, four samples from two cases of ovarian squamous cell carcinoma arising in mature cystic teratoma were also selected. Details of these cases (patients 7 and 11) have been published previously [19]. All paraffin blocks were sectioned at 3 μm on positively charged microscope slides.

#### 4.2.2 Fluorescent in situ hybridisation

FISH was performed on 3 μm tissue sections on positively charged slides using probe cocktail composed of hTERC (3q26), C-MYC (8q24) and SE 7 Triple colour (KBI-10704, Leica Microsystem). Tissue digestions and probes hybridisation was performed according to manufacture’s recommendations using Poseidon Tissue Digestion Kit I (KBI-60007 Tissue Digestion Kit I, Leica Microsystem) with the following modifications: tissue was pre-treated in Solution A (LK-110B) at 96°C to 98°C for 10 min and digested using pepsin solution (LK-110B) for 5 min. FISH digital images were captured by Nikon Eclipse fluorescence inverted microscope equipped with a charge-coupled device camera (Andor Neo sCMOS), using filter sets for DAPI/ YGFP/TRITC/CY GFP with an objective lens (Plan Apo VC 100x, Nikon). All images were captured with 100× magnification of the objective and a pixel size of 0.07 μm. For each selected field, 21 Z sections were taken with a step size of 0.3 μm. Large images of 7 × 7 fields were automatically captured from each tissue section and the 5 best fields of view with adequate tumour tissue, free of optical artefact, were chosen for further analysis with the exception of JBLAB-178 where only 2 fields of view were suitable.

#### 4.2.3 Image processing

FISH of tissue sections are noisy and display a number of recurring artefacts which can be mitigated using image preprocessing methods. The main artefacts are: bright error spots outside the nucleus which reduce true spot signal; precipitation which causes faint, erroneous spots within the nucleus; autofluorescence of areas outside the nucleus. To overcome these issues the following procedure was followed and forms part of FrenchFISH’s image prepocessing:

1. Using Fiji:

- for each field of view the position in the z-stack with the best focus was detected using Vollath’s F_4_ measure [20].
- The 4 stacks below and 5 stacks above were retained.
- A Max Intensity projection was taken across the stacks to generate a single image for further processing.
- The contrast of each spot channel was normalised and adjusted, allowing a saturation of up to 40% of the image. This allowed the weaker spot signals to be matched to the stronger, extranuclear noise spots.
2. Using R:

- Nuclear staining is segmented using the FISHalyzer package.
- Spot channel images are masked using nuclear segmentation.
- The image is filtered and normalised retaining on the top 10% of signal intensity to remove remaining autofluorescence.
- A two stage Gaussian blurring and automatic thresholding approach is applied using the Intermodes [21] method for channels with precipitation signal, and Renyi Entropy [22] method for those without precipitation, found in the autothresholdr package [23]. This combines and removes any small spot artefacts.
- A size based filter is applied for final spot segmentation.

All image processing scripts and analysis can be found in the FrenchFISH repository https://bitbucket.org/britroc/frenchfish.

#### 4.2.4 Manual spot counting

Manual spot counting was performed using IMARIS8 software following this procedure:

- Import nd2 image (21 z-stacks)
- Display in 3D
- Display DAPI channel, and switch off all other channels
- Print image
- Identify nuclei suitable for manual spot counting (none/minimal nuclear overlap cell nuclei for signal counting), circle them on the 2D image on the paper and give them numbers. Move the 3D image around to see if the nuclei are nicely separated.
- Set up aqua channel so artefacts are removed and dots clearly visible
- Set up red channel so artefacts are removed and dots clearly visible
- Set up green channel so artefacts are removed and dots clearly visible
- For every selected nuclei perform:

– Measure the size of the nuclei (x and y plane diameter)
– Count spots in aqua channel and record
– Count spots in red channel and record
– Count spots in green channel and record

## 5 Acknowledgements

We would like to acknowledge the support of The University of Cambridge, Cancer Research UK and Hutchison Whampoa Limited. Parts of this work were funded by CRUK core grant C14303/A17197 as well as A19274 (FM) and A18072 (JDB/IAMcN). We would like thank Histopathology, Biorepository, Light Microscopy Core Facilities (CRUK Cambridge Institute) and Human Research Tissue Bank and Biomedical Research Centre for study support. The Human Research Tissue Bank is supported by the NIHR Cambridge Biomedical Research Centre. We also acknowledge support from the NHS Greater Glasgow and Clyde Biorepository.

